# Reproductive sexual dimorphisms in two willow species, *Salix exigua* Nutt. *and S. nigra* Marshall

**DOI:** 10.1101/2023.05.18.541315

**Authors:** Nan Hu, Haley Hale, Brian Sanderson, Guanqiao Feng, Minghao Guo, Diksha Gambhir, Matt Olson

## Abstract

**Premise of the Research:** The prevalence of sexual dimorphisms, which evolve due to contrasting strategies to maximize reproductive success in males and females, is variable among dioecious plant species. In the *Salicaceae*, many traits have been assessed across many studies, but direct or indirect associations between these traits and reproductive allocation are often neglected. Given the dynamic evolution of sex determination systems and the strong interest in wood production in the family, we wondered whether sexual dimorphisms related to reproduction may have gone unreported. Here, we assess sexual dimorphism in reproductive traits in two species of *Salix*. Recognition of reproductive sexually dimorphic traits will contribute to understanding the evolution of sex determination systems in the Salicaceae.

**Methodology:** We conducted observational studies in natural populations to assess the presence of sexual dimorphisms in early spring bud density, catkin number, and flower number per catkin across four sampling periods in *Salix exigua*. We also analyzed flower number and catkin number per flower in *Salix nigra*.

**Pivotal Results:** We observed no sexual dimorphism in pre-season buds per branch in *S. exigua* but did find that males produced more flowers per catkin and more catkins per branch than females in both *S. exigua* and *S. nigra*.

**Conclusions:** Higher flower numbers in males compared to females is consistent with expectations from intra-sexual selection among males. The presence of reproductive sexual dimorphisms in *Salix* suggests that sexual selection may affect the evolution of mating strategies in *Salix* species, and the evolution of the sex determination system within this genus.

## Introduction

Among dioecious plant species, sexual dimorphisms (traits differing between male and female individuals) have been identified in a variety of morphological, physiological, ecological, and biochemical traits including growth and size, flower number and size, flowering duration, secondary chemistry, seed germination and dormancy, wood properties, and life history (Ashman, 2009; Barrett & Hough, 2013; Carroll & Delph, 1996; Charlesworth, 2018; Eckhart, 1999; Liu et al., 2021; Nicotra, 1999; Purrington & Schmitt, 1995; Stehlik & Barrett, 2005). These sexual dimorphisms can be categorized in multiple ways. Primary sexual dimorphisms describe differences directly associated with organ development in the androecium and gynoecium, whereas secondary sexual dimorphisms refer to characteristics beyond the differences in sexual organs.

Secondary sexual dimorphisms can be further classified into reproductive dimorphisms (traits related to gamete production, mating, and seed and fruit production) and non-reproductive dimorphisms (traits associated with all other phenotypes). For example, reproductive sexual dimorphisms have been identified in petal size, flower fragrances, and flower number (Ashman, 2009; Brandt et al., 2020; Carroll & Delph, 1996; Delph et al., 2005; Delph & Herlihy, 2012), and non-reproductive sexual dimorphisms have been found in vegetative growth and stress tolerance (Chen et al., 2010; Delph, 2019; Han et al., 2013; Olano et al., 2017; Sakai & Burris, 1985). Reproductive sexual dimorphisms are generally thought to result from natural selection that directly influences mating patterns (including sexual selection) or reproductive output (fecundity selection), whereas non-reproductive sexual dimorphisms result from different allocation tradeoffs between the sexes that ultimately lead to maximizing lifetime fitness (Liu et al., 2021; Tonnabel et al., 2021; Yang et al., 2015).

The Salicaceae, which includes both *Populus* and *Salix*, has a long history of investigations into sexual dimorphisms, dating back to seminal studies that showed habitat partitioning among arctic willows (Dawson & Bliss, 1989; Table 1). Characteristics of this family are well suited to comparative studies of sexual dimorphisms. First, all species within Salicaceae are dioecious, providing opportunities for assessing phylogenetic patterns of commonalities across species and genera; few other angiosperm families have many dioecious species (Käfer et al., 2017). Secondly, the movement of the sex chromosomes is quite active in Salicaceae (Chen et al., 2016; Hou et al., 2015; Kersten et al., 2014; Pakull et al., 2009, 2011, 2015; Pucholt et al., 2015; Sanderson et al., 2021; Yin et al., 2008; Zhou et al., 2018) and sexual dimorphisms are thought to play a part in this movement in many other systems (Charlesworth, 2018; Dean & Mank, 2014; Rice, 1984; Ross et al., 2009). Despite observations of sexual dimorphisms in many dioecious species (Delph, 2005; Lloyd & Webb, 1977), the evidence of sexual dimorphisms in the Salicaeae is mixed, and few dimorphisms appear common across species. For example, two well-replicated studies in *Populus*, found no sexual dimorphisms across 112 traits (Robinson et al., 2014) (McKown et al., 2017), whereas a recent study in Salix purpurea identified sexual dimorphisms in ∼1/2 of the measured traits (13/35; Gouker et al. 2021).

**Table 1.**
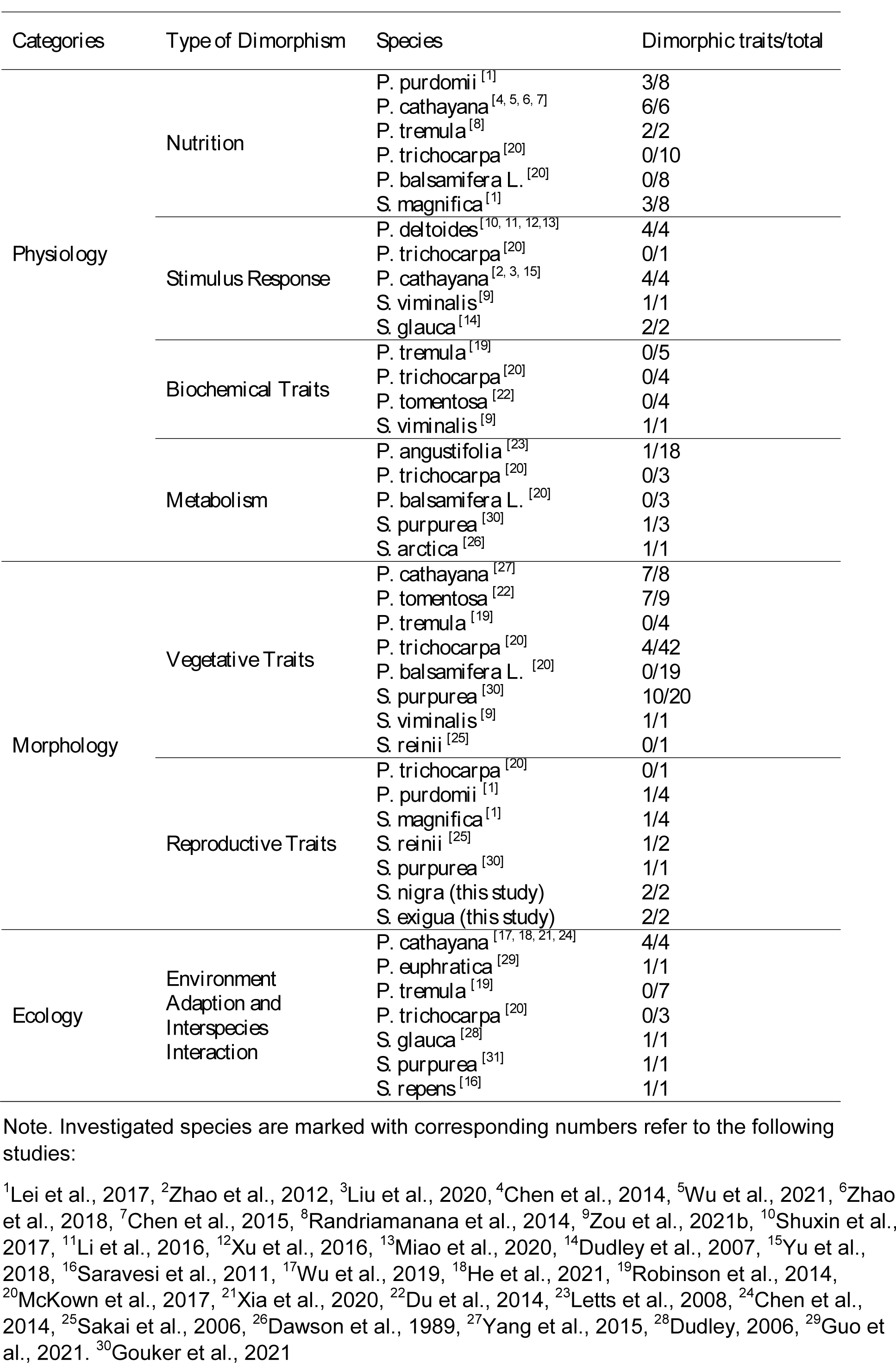
Published studies on sexual dimorphisms in *Salicaceae* species in different categories

Here we focus on natural populations of *Salix exigua* and *Salix nigra* to assess whether they exhibit sexual dimorphism in key reproductive traits. These two species represent two different subgenera, *Longifoliae* and *Protitea*, respectively, and they have sex chromosomes in different chromosomal locations, with the *S. nigra* sex chromosomes located on chromosome 7 (Sanderson et al., 2021) and the *S. exigua* sex chromosome located on chromosome 15 (Hu et al. 2022). Specifically, we ask the following questions: (1) Do females have more vegetative buds than males during early spring for *S. exigua*? (2) Are flower and catkin number sexually dimorphic for both species? We interpret our results in relation to 30 additional well-replicated studies in the Salicaceae to assess whether reproductive traits may be more likely to exhibit sexual dimorphism than non-reproductive traits and whether the extent of sexual dimorphisms may differ in *Populus* and *Salix*.

## Materials and Methods

*Salix exigua* is a dioecious shrub in subgenus *Longifoliae* (Argus, 2010) that is distributed throughout the western USA. Most flowering occurs between early April and late May, but rare catkins are produced into the fall. To assess sexual dimorphism of *S. exigua*, we sampled a linear transect next to the Rio Bonito in Fort Stanton-Snowy River Cave National Conservation Area, Lincoln County, New Mexico (33°31’40.3 “N 105°26’22.6” W). As *S. exigua* is highly clonal (Douhovnikoff & Dodd, 2003; Krasny et al., 1988), we sampled neighboring ramets only if they were different sexes, so they were clearly different genets, or if they were spaced at intervals of at least 10 meters. Individual sexes were obtained by observing flower structures during flowethe ring season. Specifically, males were identified as individuals with anthers and no carpels; females were identified as individuals with carpels and no anthers.

*Salix nigra* is a tree-form willow in subgenus *Protitea* that is distributed across the central and eastern USA (Argus, 2010). We sampled a population on the western edge of its range in Dickens Springs County Park, Dickens Springs, Texas (33°37’44.0 “N 100°49’36.7”W). O ver 50 trees were tagged and sexed in this population from an earlier experiment designed to map the sex chromosome (Sanderson et al., 2021). Based on our previous sampling, we determined that *Salix nigra* flowers from late April to early May and exhibits much less clonal reproduction than *S. exigua*.

Sexual dimorphism of three reproductive traits was investigated in *S. exigua*: spring bud density prior to flowering, the density of catkins on stems, and flower numbers per catkin. *Salix exigua* exhibits an extended and intermittent flowering period and males quickly drop their catkins after anthesis. Thus, we estimated total catkin production using multiple observations across the flowering period. Prior to the flowering season in March 2018, one similar-sized branch was randomly selected, tagged, and marked on 64 random individuals. Individuals were at least 10 m apart to minimize sampling from the same genotype; subsequent genotyping indicated that this spacing was sufficient (Hu, unpublished data). Bud density on sampled branches was estimated prior to flowering by counting the number of buds on the last 40 cm of each branch. Bud type is undifferentiated in *S. exigua* in early spring but can be assessed by tracing catkin development. Marked plants were revisited every two weeks to count the number of catkins in anthesis (males) or receptive (females) on the marked portion of the stems. Because our sample sizes were small and many *S. exigua* individuals did not flower at all sampling times, analyses using repeated measures models did not converge.

Nonetheless, our observations indicated that total catkin production could be estimated by summing flowering catkins across all four sampling periods because the same catkin was not flowering on multiple sampling periods. For 8 males and 12 females, no flowers were in anthesis (males) or receptive (females) during the sampling periods before and after catkins were flowering on the labeled branch. Finally, in mid-May, flower numbers per catkin and the length of the catkin were recorded for 3 randomly selected catkins for 8 males and 8 females. Flower number per catkin was measured on different individuals than those on which we measured the other traits.

Flowering phenology in *S. nigra* is much more synchronous than in *S. exigua*, allowing us to estimate catkin production by sampling at only one time period. For *S. nigra,* two traits were investigated: the number of catkins per branch and flower numbers per catkin. To estimate the number of catkins per branch, we sampled 3 randomly selected branches on previously tagged male (n=11) and female (n=9) trees on branches that could be reached from the ground with a pole saw. The number of total twigs from each sampled branch, the number of flowering and non-flowering twigs, and the diameters of sampled branches also were recorded. The numbers of flowers per catkin and catkin lengths were measured on 3 randomly chosen catkins from 5 males and 5 females. Catkin numbers and flowers per catkin were measured on different individuals.

We used Student’s t-test in R v.4.0.1 to test for significant trait differences between males and females (R Core Team, 2021). Plotting was done using the ggplot2 package (Wickham, 2016).

## Results

After flowering, we identified tagged individuals of *Salix exigua* to be 21 males, 25 females, and 18 of unknown sex. This sex ratio was not significantly different from 1:1 (χ^2^ =0.348, P > 0.05). Males and females had similar numbers of pre-season buds per branch on *S. exigua* (Fig. 1, p = 0.09). Both *S. exigua* and *S. nigra* males produced more catkins than females across the flowering season (Fig 2A, p < 0.05; Fig. 2B, p < 0.001). Males also produced more flowers per catkin than females in both species (Fig. 2A; Fig 2B, p < 0.001). Catkin lengths did not differ between males and females in either *S. exigua* or *S. nigra* (P = 0.19; P = 0.24).

**Figure 1.**
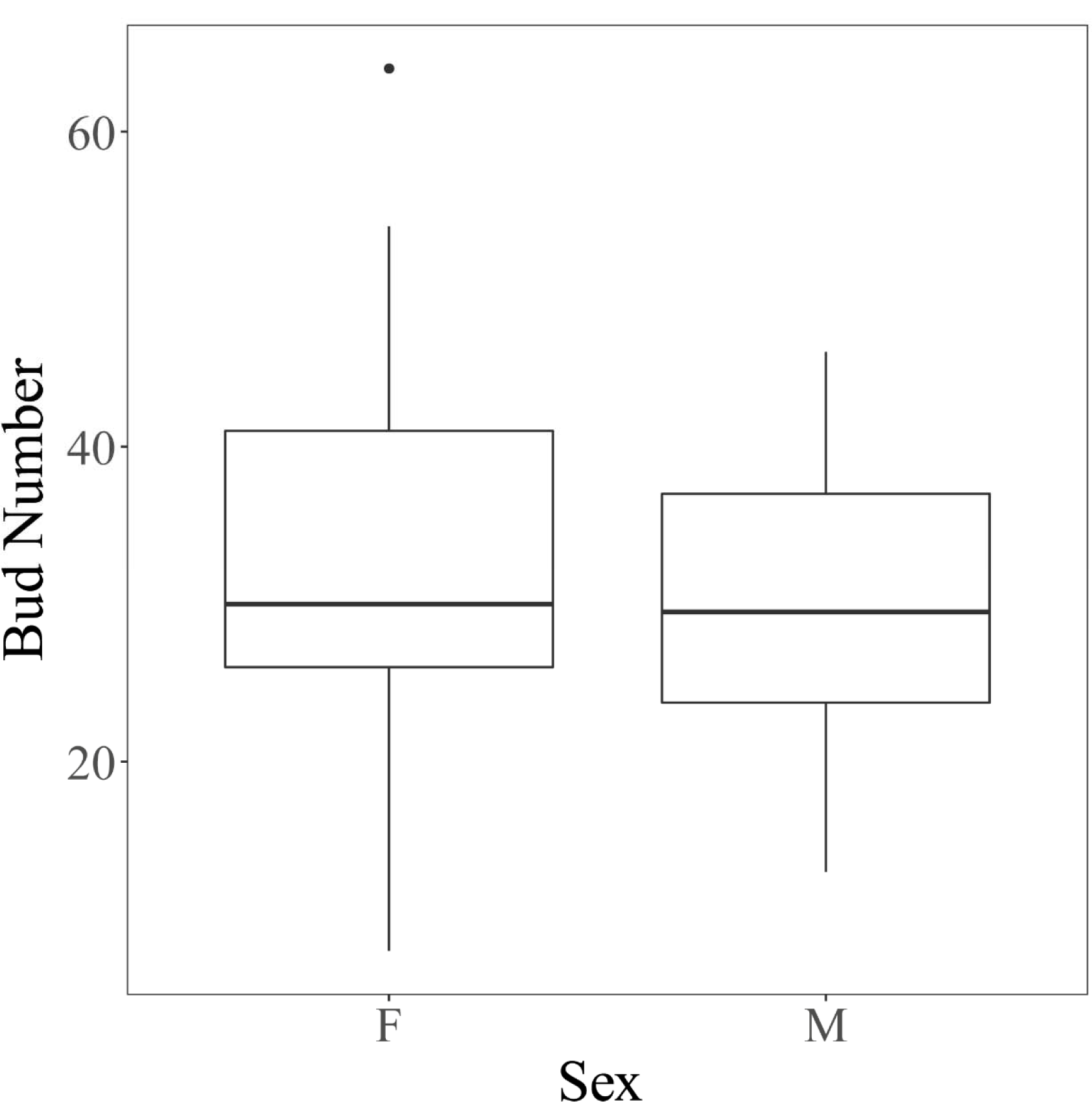
Boxplot of spring bud numbers (both vegetative and floral) per sampled branch for males and females in S. exigua. F: Female. M: Male. No sexual dimorphism was detected (P = 0.08).

**Figure 2.**
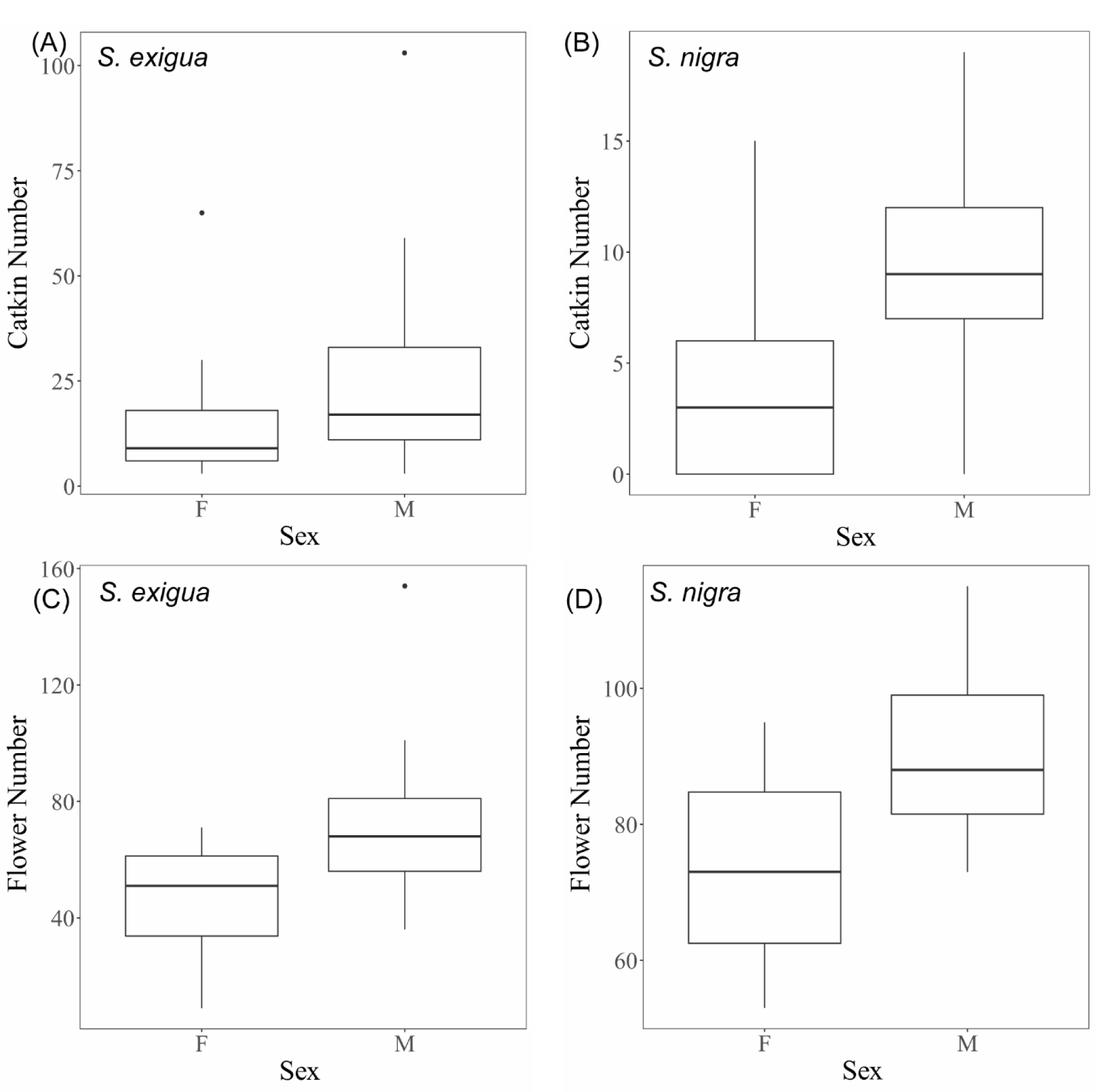
Boxplots of sexual dimorphisms in S. exigua and S. nigra. F: Female. M: Male. All traits in both species are significant (P < 0.05). (A) Catkin number for males and females in S. exigua. (B) Catkin number for males and females in S. nigra. (C) Flower number per catkin in S. exigua. (D) Flower number per catkin in S. nigra.

Three traits were measured as potential covariates for bud number and catkin length in *S. exigua* and one was measured in *S. nigra*. None of them accounted for a significant proportion of variance and were dropped as covariates, but we document them for completeness. None of the vegetative measurements including stem length, diameter, and total twig length were sexually dimorphic in *S. exigua* (P = 0.52; P = 0.09; P = 0.73), and stem length was not sexually dimorphic in *S. nigra* (P = 0.23) (Table S1).

## Discussion

We identified two reproductive sexual dimorphisms in both *S. exigua* and *S. nigra* showing that males produced more catkins and more flowers per catkin than females; however, we found no differences in the numbers of preseason buds or stem and twig sizes produced by males and females of *S. exigua*. These patterns are consistent with expectations when intra-sex selection among males for access to mates increases flower or pollen production (Bateman, 1948; Minnaar et al., 2019; Muchhala et al., 2010) and suggest that some willows may have evolved sexual dimorphisms in response to selective pressures to maximize mating opportunities. Notably, although bud numbers did not differ for males and females of *S. exigua* (preseason bud numbers were not counted in *S. nigra*), catkin numbers were greater for males indicating that females produced more vegetative bud than males. Unlike in Populus, floral and vegetative buds are indistinguishable in *S. exigua*.

Our discoveries contribute to understanding secondary reproductive dimorphisms in the Salicaceae family. Table 1 shows the results of our review of 31 studies (including ours) across 20 species in the Salicaceae with sample sizes of at least 20 male and 20 female genotypes. Before the current study, only four studies on five species included traits that related to sexual reproduction and mating, and only two studies focused on a willow species. Of five species studied for reproductive dimorphisms, four identified at least one sexual dimorphism (Table 1). In other angiosperm systems such as *Silene*, *Rumex*, and *Leucadendron*, sexual dimorphisms in petal size, flower shape, flower color, flower number, and other features influencing mating have been the subjects of intensive investigations (Delph et al., 2005; Harris & Pannell, 2010; Midgley, 2010; Stehlik & Barrett, 2005; Zemp et al., 2018). These features often experience selection for pollinator attraction which drives the differentiation of strategies between males and females (Kiester et al., 1984). This is in contrast with studies in the *Salicaceae* (Table 1), which have focused more on wood properties, stress response, and leaf traits. This bias is likely derived from both the early studies of *Salix arctica* that found ecological and physiological differences between males and females (Dawson & Bliss, 1989) and from the economic potential of salicaceous plants related to biomass yield (Rowe et al., 2013; Sevel et al., 2012; Tullus et al., 2012). We propose that studies focused on sexual dimorphisms in reproductive traits would have stronger trait differences than those focused on non-reproductive traits and would be useful for understanding patterns of evolution of sexual systems in other Salicaceous species. Moreover, considering the recent advances in understanding the evolution of sex chromosomes in the *Salicaceae* (Cronk & Müller, 2020; Sanderson et al., 2021; Wei et al., 2020; Xue et al., 2020; W. Yang et al., 2021; Zhou et al., 2018a), additional studies of mating patterns and their relationships with sexual dimorphisms are warranted.

Sexual dimorphisms identified in *S. exigua* and *S. nigra* represent fundamentally different resource allocation strategies between the sexes. Based on these differences, we hypothesize that males benefit from producing more flowers and catkins that both attract more pollinators to outcompete other males for access to mates and produce more pollen to increase the potential for increased numbers of mates. Male catkins in Salicaceae are often considered showier than females because they are larger with more flowers (Cronk et al., 2015), and the anthers are bright yellow and extend farther from the base of the flower than the pistil, making the catkin have a larger axis (Cronk et al., 2015). Increased catkin and flower numbers have been shown in other species to attract more pollinators and increase male mating success (Matsuhisa & Ushimaru, 2015; Yakimowski et al., 2011). More conspicuous male flowers also have been shown to drive pollinator movement by encouraging pollinators to visit male flowers before visiting female flowers, as in *Eurya japonica* (Tsuji et al., 2020). Recent studies on *S. nigra* showed that insect communities have higher diversity in male than in female flowers (Simon et al., 2021), which may indicate higher attraction to pollinators in males.

We showed that the number of flowers per catkin and the number of catkins per branch is shared sexual dimorphisms between these two species. Considering that these two species differ in the genomic locations of their sex determination genes, it is worth considering whether this similarity evolved twice independently or if these dimorphisms have been maintained for long time periods and inherited from a shared ancestor. The ultimate genetic control of sexual dimorphisms will always map to the sex chromosome, although developmental genes in the pathway hierarchy just below the sex determination genes on the sex chromosome will usually map to autosomes (Feng et al., 2020). The genetic factors controlling the sexually dimorphic traits in these two *Salix* species are currently unknown but are either pleiotropic effects of the sex determination genes, are controlled by the same accessory genes located in the non-recombining sex determination region, or are controlled by different independently-evolved accessory genes located in the non-recombining sex determination region.

Wang et al. (2022) showed that only one protein-coding gene is shared between sex chromosome 7 in *S. chaemeneloides* and sex chromosome 15 in *S. arbutifolia*, two species that are closely related to and share sex determination regions with *S. nigra* and *S. exigua*, respectively. However, this gene is not related to flower development in *A. thaliana* (AT4G23160) and is thus unlikely to function as a gene controlling these sexual dimorphisms. Assuming that there are no shared genes influencing sexual dimorphisms in catkin production either the sex determination gene has maintained the same pleiotropic effects since the divergence of *S. nigra* and *S. exigua* or the sexual dimorphism evolved independently in both species. Understanding the genetic basis of these floral sexual dimorphisms and their effects on fitness would aid in testing whether the sexually antagonistic selection hypothesis drove the movement of the sex chromosome in ancestors of these two species (Rice, 1987; Saunders et al., 2019; van Doorn & Kirkpatrick, 2007, 2010). The sexual antagonism hypothesis, however, not only proposes that sexually antagonistic genes are located on the sex chromosomes, but also that a new gene influencing a sexually antagonistic trait arises on an autosome and induces the movement. Future research to understand the effects of the accessory genes on the non-recombining portions of the sex chromosomes in *S. nigra* and *S. exigua* would aid in testing this hypothesis as a mechanism driving the movement of sex chromosomes in plants.

## Acknowledgment

We greatly thank Lei Chen from Sichuan University, China, inspired us in statistical analysis. We also thank Ashmita Khanal for giving advice on figures. N. Hu and M. Olson designed the experiment. H. Hale selected sampling populations as well as performed a preliminary study. All the authors were involved in data sampling and fieldwork. N. Hu, G. Feng, B. Sanderson, and M. Olson collected *S. exigua* data. N. Hu, M. Olson, M. Guo, D. Gambhir collected *S. nigra* data. B. Sanderson and G. Feng offered detailed feedback on the manuscript. N. Hu and M. Olson performed the data analysis and wrote the manuscript. The manuscript was approved by all the authors before submission. This work was supported by the National Science Foundation under the award IOS-1542599.

## Notes

### Competing Interest Statement

The authors have declared no competing interest.

