## Supplemental Table 1 for "Reproductive sexual dimorphisms in two willow species, *Salix exigua* Nutt. *and S. nigra* Marshall"

Tables 1a-g: Type II analysis of variance tables for numbers of pre-season buds per branch on *Salix exigua* and catkins per stem, numbers of flowers per catkin, and catkin length on both *S. exigua* and *S. nigra*. Satterthwaite’s method was used for calculating F statistics for random effects in mixed models.

a) Response: *Salix exigua* pre-season buds per branch

|  | Df | Sum-Sq | Mean-Sq | F-value | Pr(>F) |
| --- | --- | --- | --- | --- | --- |
| Sex | 1 | 180.1 | 180.1 | 2.3358 | 0.1747 |
| Diameter | 1 | 136.2 | 136.2 | 1.7664 | 0.1838 |

b) Response: *Salix exigua* catkins per stem

|  | Df | Sum-Sq | Mean-Sq | F-value | Pr(>F) |
| --- | --- | --- | --- | --- | --- |
| Sex | 1 | 2.9896 | 2.98962 | 4.6032 | 0.0376* |
| stemLength | 1 | 0.1718 | 0.17180 | 0.2645 | 0.6097 |

c) Response: *Salix nigra* catkins per stem

|  | Sum-Sq | Mean-Sq | NumDF | DenDF | F-value | Pr(>F) |
| --- | --- | --- | --- | --- | --- | --- |
| Sex | 67.175 | 67.175 | 1 | 18.603 | 6.9386 | 0.0165* |
| stemLength | 55.18 | 55.18 | 1 | 54.767 | 5.6997 | 0.0205* |

d) Response: *Salix exigua* flowers per catkin

|  | Sum-Sq | Mean-Sq | NumDF | DenDF | F-value | Pr(>F) |
| --- | --- | --- | --- | --- | --- | --- |
| Sex | 2071.5 | 2071.5 | 1 | 14 | 8.3032 | 0.0121* |

e) Response: *Salix nigra* flowers per catkin

|  | Sum-Sq | Mean-Sq | NumDF | DenDF | F-value | Pr(>F) |
| --- | --- | --- | --- | --- | --- | --- |
| Sex | 389.9 | 389.9 | 1 | 9 | 6.0012 | 0.03677* |

f) Response: *Salix exigua* catkin length

|  | Sum-Sq | Mean-Sq | NumDF | DenDF | F-value | Pr(>F) |
| --- | --- | --- | --- | --- | --- | --- |
| Sex | 25.957 | 25.957 | 1 | 14 | 1.8596 | 0.1942 |

g) Response: *Salix nigra* catkin length

|  | Sum-Sq | Mean-Sq | NumDF | DenDF | F-value | Pr(>F) |
| --- | --- | --- | --- | --- | --- | --- |
| Sex | 60.364 | 60.364 | 1 | 9 | 1.5575 | 0.2435 |

Note. Df: Degree of freedom. Sum-Sq: Sum of squared value. Mean-Sq: Mean of Squared value. Pr: Probability. NumDF: Nominator degree of freedom. DenDF: Denominator degree of freedom.

*: significance level at p < 0.05
